# Comprehensive genome sequencing analysis as a promising option in the prenatal diagnosis of fetal structural anomalies: a prospective study

**DOI:** 10.1101/2020.08.22.260893

**Authors:** Jia Zhou, Ziying Yang, Jun Sun, Lipei Liu, Xinyao Zhou, Fengxia Liu, Ya Xing, Shuge Cui, Shiyi Xiong, Xiaoyu Liu, Yingjun Yang, Xiuxiu Wei, Gang Zou, Zhonghua Wang, Xing Wei, Yaoshen Wang, Yun Zhang, Saiying Yan, Fengyu Wu, Fanwei Zeng, Tao Duan, Jian Wang, Yaping Yang, Zhiyu Peng, Luming Sun

**Affiliations:** Shanghai First Maternity and Infant Hospital, Tongji University School of Medicine, Shanghai, China; BGI Genomics, BGI-Shenzhen, Shenzhen, China; Tianjin Medical Laboratory, BGI-Tianjin, BGI-Shenzhen, Tianjin, China; Department of Medical Genetics and Molecular Diagnostic Laboratory, Shanghai Children’s Medical Center, Shanghai Jiaotong University School of Medicine; Department of Biology, Faculty of Science, University of Copenhagen, Copenhagen, DK-2200, Denmark; AiLife Diagnostics, Pearland, TX 77584, USA

**Keywords:** Genome sequencing, Chromosomal microarray analysis, Exome sequencing, Fetal structural or growth anomalies, Prenatal diagnosis

## Abstract

**Purpose:** Genome sequencing (GS) is a powerful tool for postnatal genetic diagnosis, but relevant clinical studies in the field of prenatal diagnosis are few. We aimed to evaluate the feasibility of GS as a first-line approach in prenatal diagnosis and compare its clinical value with the chromosomal microarray analysis (CMA) plus exome sequencing (ES) sequential testing.

**Methods:** We applied trio GS (∼40-fold) in parallel with CMA plus ES to investigate the genetic basis for structural or growth anomalies in 111 fetuses and compared their results.

**Results:** GS covered all genetic variants in 22 diagnosed cases detected by CMA plus ES, yielding a diagnostic rate of 19.8% (22/110). Moreover, GS provided more comprehensive and precise genetic information than CMA plus ES, revealing twin fetuses with an imbalanced translocation arising from a balanced paternal translocation and one fetus with an extra pathogenic variant in the *GJA8* gene, and incidentally identified intrauterine CMV infection in a growth-restricted fetus.

**Conclusion:** Compared with CMA plus ES, GS offers a more comprehensive view of the genetic etiology of fetal anomalies and provides clues for nongenetic factors such as intrauterine infection. Our study demonstrates the feasibility of GS as a promising first-line test in prenatal diagnosis.

## INTRODUCTION

Congenital fetal anomalies occur in approximately 3% of pregnancies[1], and many of these anomalies have an underlying genetic etiology. Identification of the genetic basis of the anomalies enables informed decision-making, improves perinatal care, and helps to assess recurrence risk for future pregnancies. Chromosomal microarray analysis (CMA) has been broadly adopted to detect copy number variants (CNVs) in prenatal diagnoses with an additional 6% of diagnostic yield over standard karyotyping in fetuses with structural anomalies observed by ultrasound[2, 3]. CMA can detect CNVs as small as 10-100 kb in length depending on the probe density. However, smaller variants, such as single nucleotide variants (SNVs) and small insertions or deletions (INDELs), which also contribute to a substantial portion of genetic disorders remain undetectable by this approach[4-7]. Exome sequencing (ES), which detects SNVs, INDELs, and CNVs covering multiple exons, has been proven to be a powerful tool in prenatal diagnosis. In clinical practice, ES can be conducted in CMA-negative cases to further search for single-base lesions. Emerging studies have shown that ES has a detection rate of 8.5% to 10% in fetal structural abnormalities with normal karyotype and CMA results[8, 9].

Even though the combination of CMA and ES considerably increases the diagnostic yield[10, 11], it is a time-consuming stepwise approach and requires a large amount of DNA. Given the time-sensitive nature of the prenatal stage and the potential inaccessibility of adequate fetal samples, the clinical utility of this sequential scheme in prenatal diagnosis is undetermined. Genome sequencing (GS) is capable of detecting almost all types of genomic variants with a low input-DNA requirement (∼100 ng) and is proposed to be beneficial in prenatal diagnosis[12, 13].

Several studies have supported GS as the first-tier test in the postnatal evaluation of patients with unexplained developmental delay, intellectual disability, congenital anomalies, and autism spectrum disorder[14-16]. However, except for small case series with a limited number of highly selected fetuses, few studies have tried to evaluate the feasibility and value of GS in prenatal diagnosis. Talkowski et al.[17] identified precise translocation breakpoints that directly disrupted *CHD7* and *LMBRD1* by using ∼12-fold GS in an undiagnosed prenatal sample, and this finding could not have been reliably inferred from conventional karyotyping. Choy et al.[18] demonstrated that 30-fold GS could provide a two-fold increase in diagnostic yield (32.0%, 16/50) in fetuses with increased nuchal translucency (≥3.5 mm) compared with routine CMA and/or karyotyping (16.0%, 8/50). The additional diagnoses by GS include 7 (14%, 7/50) cases with pathogenic or likely pathogenic (P/LP) SNVs/INDELs, and their result demonstrated the potential of GS to replace the combination of CMA and ES in prenatal diagnosis.

To comprehensively investigate the diagnostic yield of GS and the feasibility of GS as a first-line approach in prenatal diagnosis, we prospectively applied GS to 111 fetuses with structural or growth anomalies and compared the results of GS with CMA plus ES.

## PATIENTS AND METHODS

### Study design and participants

The flowchart of this study is presented in Figure 1. The parents of 111 fetuses with structural or growth anomalies identified by sonographic examination and indicated for prenatal genetic tests were recruited from Shanghai First Maternity and Infant Hospital. Fetal samples were obtained through an invasive diagnostic procedure such as amniocentesis, chorionic villus sampling or cordocentesis. Parental peripheral blood samples were collected. Samples from fetuses were subjected to two test strategies: parent-fetus trio GS and CMA plus ES. ES was sequentially performed only in samples with negative CMA results. P/LP SNVs/INDELs were validated by Sanger sequencing, while P/LP CNVs identified by GS were cross-validated with the results of CMA. Quantitative PCR (qPCR) was conducted for additional P/LP CNVs identified by GS. Genetic counseling was performed both before and after the prenatal genetic tests, and pregnant women were informed about the results of the tests they had taken. Pregnancy outcomes were followed up. This study protocol was approved by the Ethics Committee of Shanghai First Maternity and Infant Hospital. Written informed consent was obtained from pregnant women and their partners.

**Figure 1.**
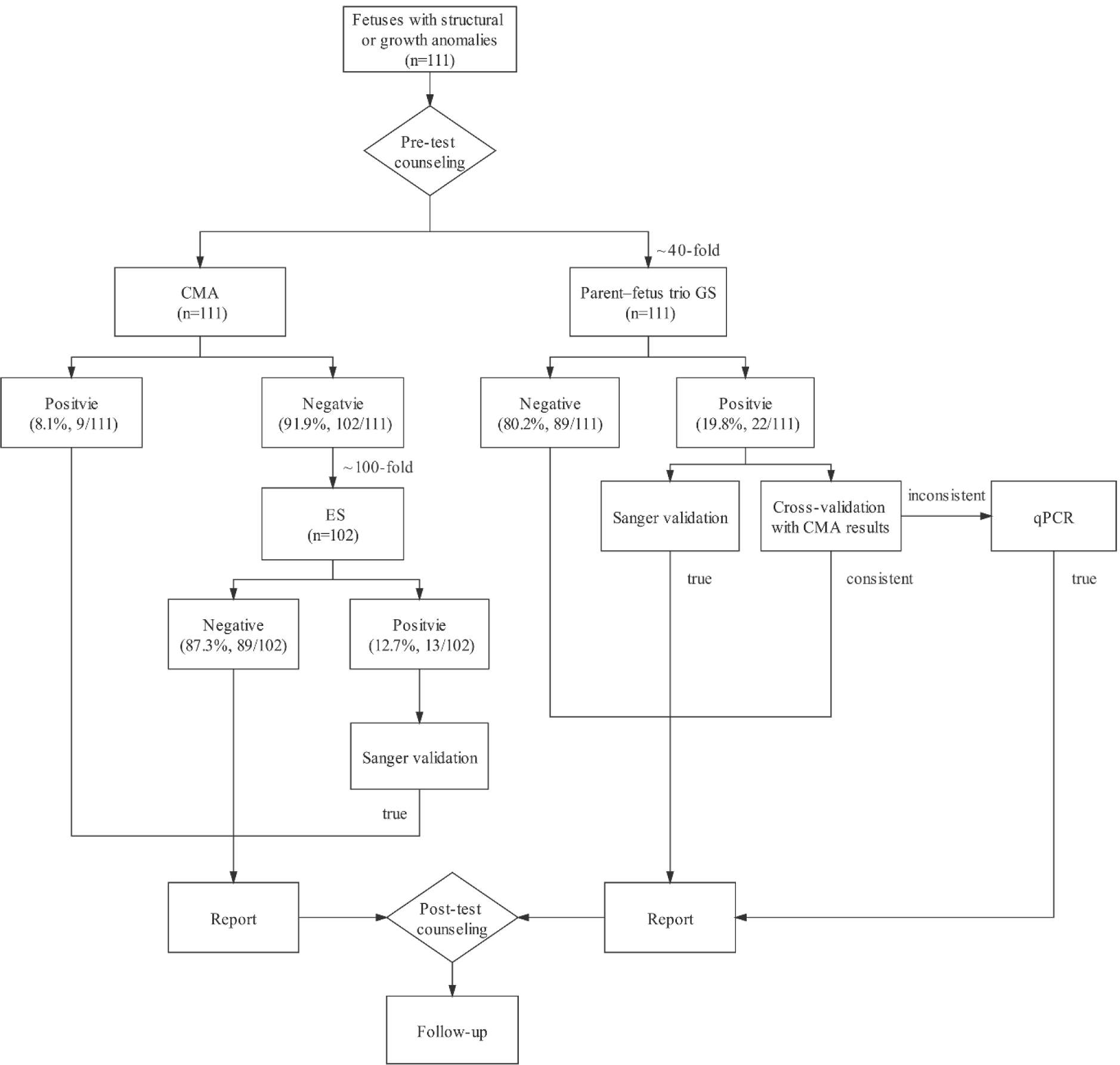
Flowchart of the study. A total of 111 fetuses with structural or growth anomalies were subjected to two test strategies: trio genome sequencing (GS) and chromosomal microarray (CMA) plus exome sequencing (ES). ES was sequentially performed in only 102 fetuses with negative CMA results. The positive/negative rates are provided in each box.

### GS

Parental and fetal samples were sequenced concurrently (fetus-parent trios or dyads testing). First, 80-200 ng of genomic DNA from each sample was sheared by the Covaris S220 Focused Ultrasonicator (Covaris, Woburn, USA). The fragmented DNA was further processed with AMPure XP beads (Life Sciences, Indianapolis, USA) to obtain 100 bp-300 bp fragments. Library construction, including end repair, A-tailing, adapter ligation, and 7 cycles of PCR amplification, was subsequently conducted. The PCR products were then heat-denatured to form single-strand DNAs, followed by circularization with DNA ligase, and the remaining linear molecule was digested with the exonuclease. After construction of the DNA nanoballs, paired-end sequencing with 100 bp at each end was carried out for each sample with a minimal read depth of 40-fold on the MGISEQ-2000 platform (MGI, Wuhan, China)[19].

Data analysis and bioinformatics processing are described in detail in Supplementary Materials and Methods. Briefly, sequencing reads were aligned to the NCBI37/hg19 assembly using the Burrows-Wheeler Aligner (BWA, version 0.7.17) with default parameters. SNVs and INDELs were called using the Genome Analysis Toolkit (GATK, version 4.0.11) and annotated by the in-house pipeline BGICG_Anno (version 0.3.9). CNVs and structural variants (SVs) were analyzed using CNVnator (version v0.3.2) and LUMPY (version 0.3.0), respectively. The outputs of these callers were merged into a single-variant call format (VCF) file per trio and annotated with public databases and our in-house databases. The results of the SNV/INDEL, CNV and SV analyses were integrated and reviewed for the interpretation of pathogenicity.

### CMA

CytoScan 750K (Affymetrix, Santa Clara, USA) was the CMA platform used in the prenatal genetic diagnosis center of Shanghai First Maternity and Infant Hospital. A total of 250 ng of DNA was required for routine prenatal CMA testing according to the manufacturer’s protocols[20]. CNVs were analyzed via CHAS 2.0 software (NCBI37/hg19), and the reporting size threshold of the CNVs was set at 100 kb with a marker count of ≥50.

### ES

ES was also performed by inputting 150-300 ng of genomic DNA from each sample and was prepared with the same library construction procedure as GS. Exome capture using the MGIEasy Exome Capture V4 Probe (MGI, Wuhan, China) was followed by paired-end read sequencing (2 × 100 bp read length) on the MGISEQ-2000 platform with an average depth of ≥100-fold. Exome sequencing data analysis was performed as previously described[21].

### Data interpretation and reporting

CNVs detected by CMA and GS were interpreted following standards of the American College of Medical Genetics and Genomics (ACMG) and the Clinical Genome Resource (ClinGen)[22]. Among the large numbers of SNVs/INDELs, we prioritized candidate causative SNVs/INDELs per the following criteria: 1) absent or with a minor allele frequency ≤1% in the databases of ExAC (http://exac.broadinstitute.org) and gnomAD (https://gnomad.broadinstitute.org), corresponding to the variant evidence as PM2; 2) family segregation information that was consistent with the inheritance of the variants (PS2/PM6/PM3); 3) supporting evidence from published literature (e.g., PS1/PS3/PS4/PM5); 4) null variants or CNVs that overlapped with established triplosensitive, haploinsufficient genes or genomic regions (PVS1); 5) conservation and predicted impact on coding and noncoding sequence (PP3); and 6) relevance to the fetal clinical phenotype (PP4). All the selected variants were assessed for pathogenicity based on the adapted ACMG guidelines and ClinGen sequence variant interpretation working group per updated recommendations for the ACMG criteria[23-26].

All of the candidate causative variants were reviewed by a multidisciplinary team of fetal medicine specialists, genetic counselors and geneticists. The result that was unanimously agreed upon to explain the fetal phenotype included P/LP variants consistent with the inheritance pattern and phenotype of related disorders, as well as variants of unknown significance (VUS) among Online Mendelian Inheritance in Man (OMIM) disease-causing genes that matched the fetal phenotype and was found in trans with a P/LP variant in an autosomal recessive condition, was designated positive or diagnostic; otherwise, was designated negative or undiagnostic. The pregnant women and their partners were informed of the ACMG secondary findings[27], and fetal and parental incidental findings were reported only when consented in the pretest informed consent process according to the ACMG document[28].

### Data validation

SNVs/INDELs/SVs were validated by Sanger sequencing. For CNV validation, qPCR was conducted for additional P/LP CNVs identified by GS. The HCMV Real-Time PCR Kit (Liferiver, Shanghai, China) was used for validation of one case that was identified by GS as suspicious for CMV infection.

## RESULTS

Between November 2019 and January 2020, 111 fetuses with structural or growth anomalies and 209 matched parental samples were eligible for inclusion in our study. Samples from a total of 320 individuals (106 fetus-parental trios, including 4 sets of twins and 5 fetus-parent dyads) were analyzed by GS, while the 111 fetal samples were also analyzed by CMA in parallel. Samples from 102 fetuses with negative CMA results were subsequently analyzed by ES. The fetuses were assessed at a median gestational age of 24 (range 16–34) weeks. The results of the Down syndrome maternal-serum screening test or noninvasive prenatal screening test were recorded when available. The detailed clinical information is available in Table S1.

### Comparison of GS and CMA plus ES

With CMA plus ES strategy, 6 (5.4%, 6/111) fetuses with aneuploidies (3 with trisomy 21, 2 with trisomy 13 and 1 with trisomy 18) and 3 (2.7%, 3/111) fetuses with P/LP CNVs were identified by CMA (Table 1), while 13 (11.7%, 13/111) fetuses with P/LP SNVs/INDELs were identified by ES (Table 2), providing a diagnostic yield of 19.8% (22/111). In comparison, GS not only detected all genetic variants in 22 cases diagnosed by CMA plus ES but also reported more types of variants and provided more comprehensive genetic information (Figure 2). Specifically, in cases 102-1 and 102-2, which were twins with severe hydroderma and pleural effusion, a 19.7-Mb deletion on the terminal q-arm of chromosome 13 (13q32.1q34), in which 3 genes were associated with autosomal dominant developmental disorder, and a 1.6-Mb duplication on the terminal q-arm of chromosome 15 (15q26.3) were both identified by CMA and GS (Figure 3A). However, GS data indicated that these two CNVs were caused by an imbalanced translocation between chromosome 13 and chromosome 15 in both twin fetuses when we checked the precise breakpoints of the detected CNVs through Integrative Genomics Viewer (IGV). This imbalanced translocation was then confirmed by Sanger sequencing (Figure S1). The fetal karyotype strongly suggests parental translocations between chromosomes 13 and 15, and indeed, a balanced translocation was identified in the father of the twins in the analysis of the GS data, while their mother’s karyotype was normal. We further delineated aberrations in the breakpoint region by using IGV and confirmed the breakpoints of the balanced interchromosomal translocation by Sanger sequencing (Figure 3B). There was a templated 13-bp duplication and a 34-bp deletion in the breakpoint junction on chromosomes 13 and 15, respectively (Figure 3C), indicating that the breakpoint repair mechanism might be caused by nonhomologous end joining with a high degree of precision but occasional deletion during DNA processing before ligation[29]. Therefore, for this family, GS provided comprehensive and precise genetic information by detecting more types of variants that might not be detectable by CMA or ES.

**Table 1.**
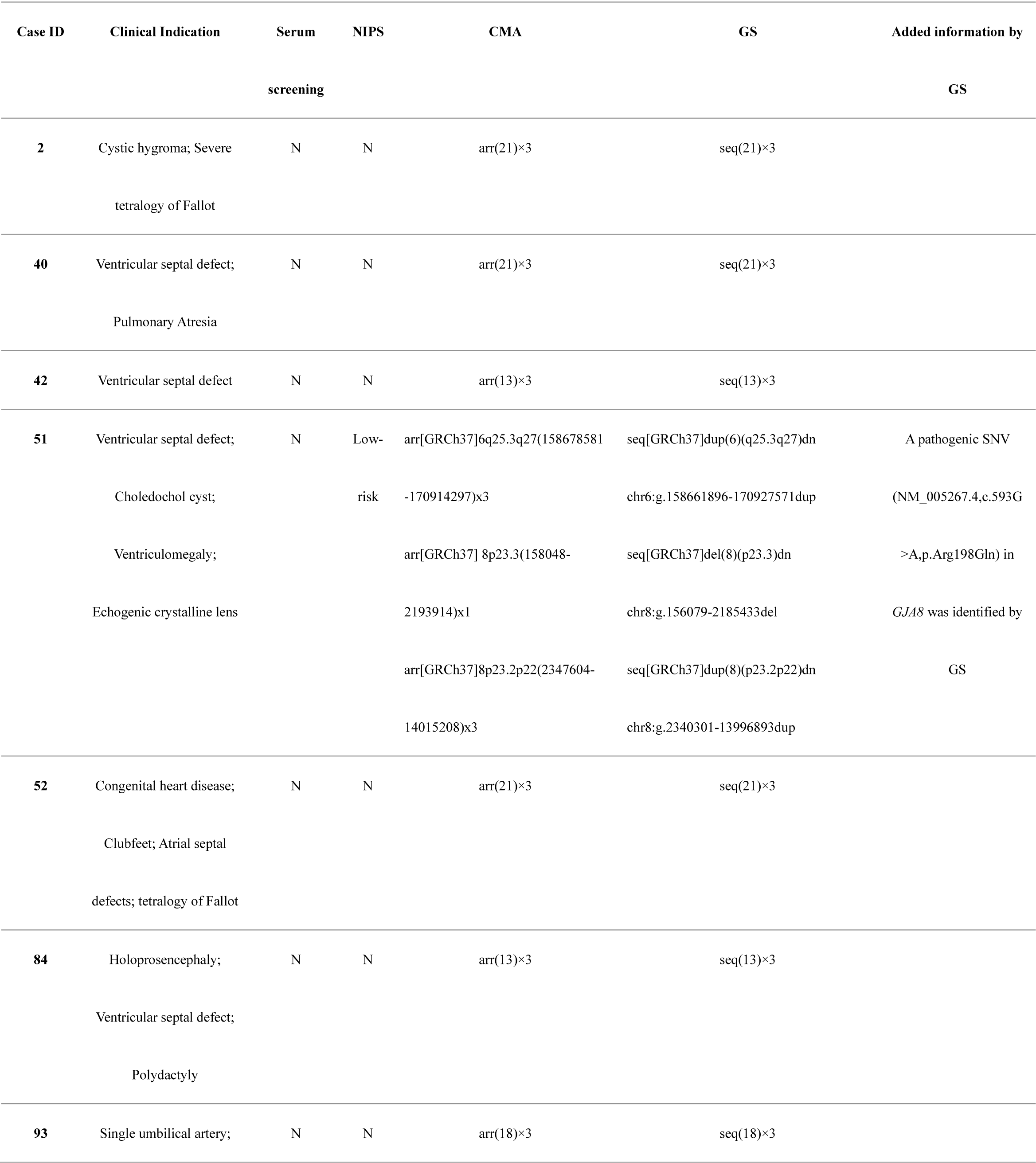
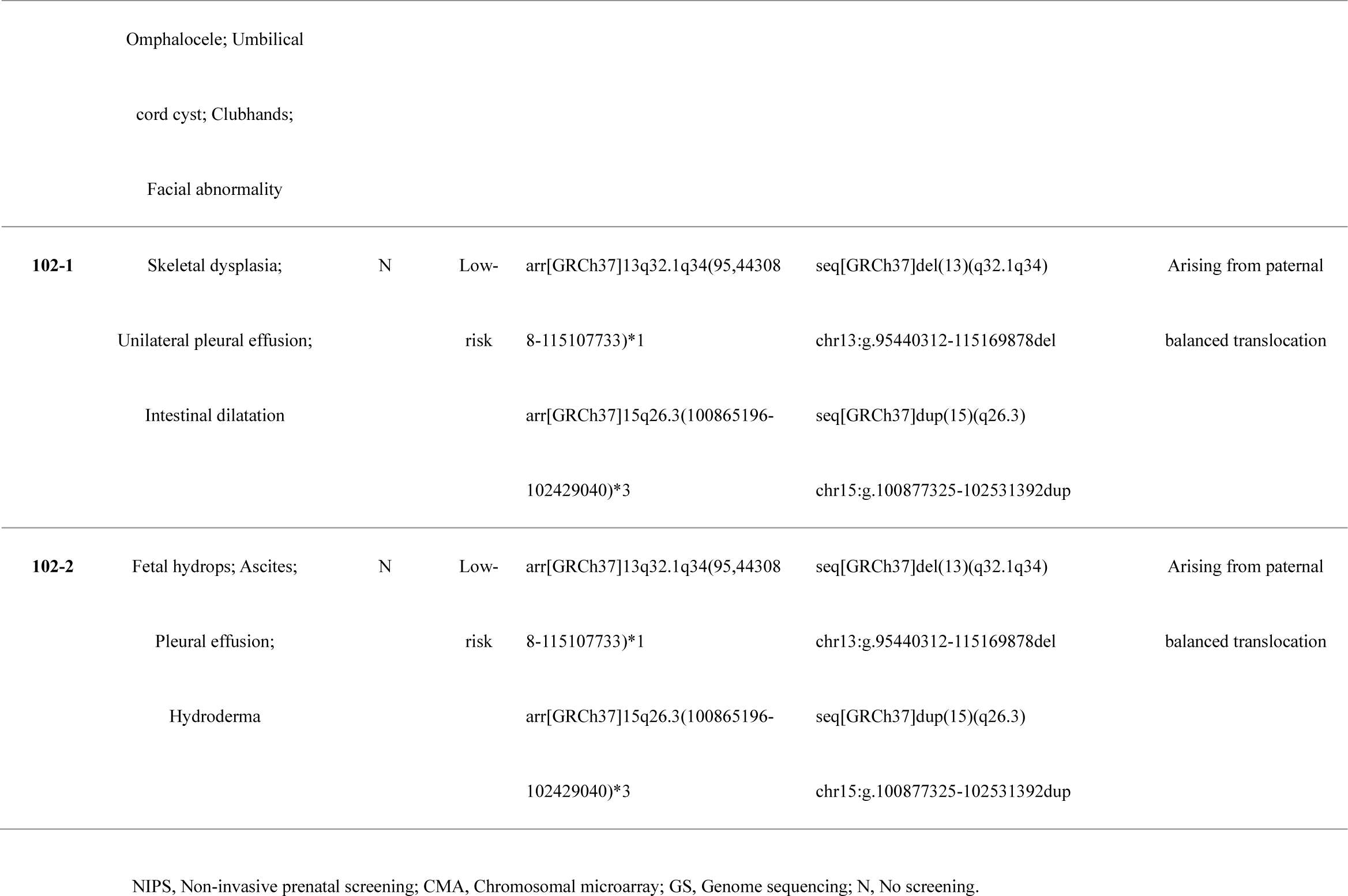
Summary of numerical disorder and pathogenic or likely pathogenic copy number variants (CNVs) detected by CMA and GS.

**Table 2.**
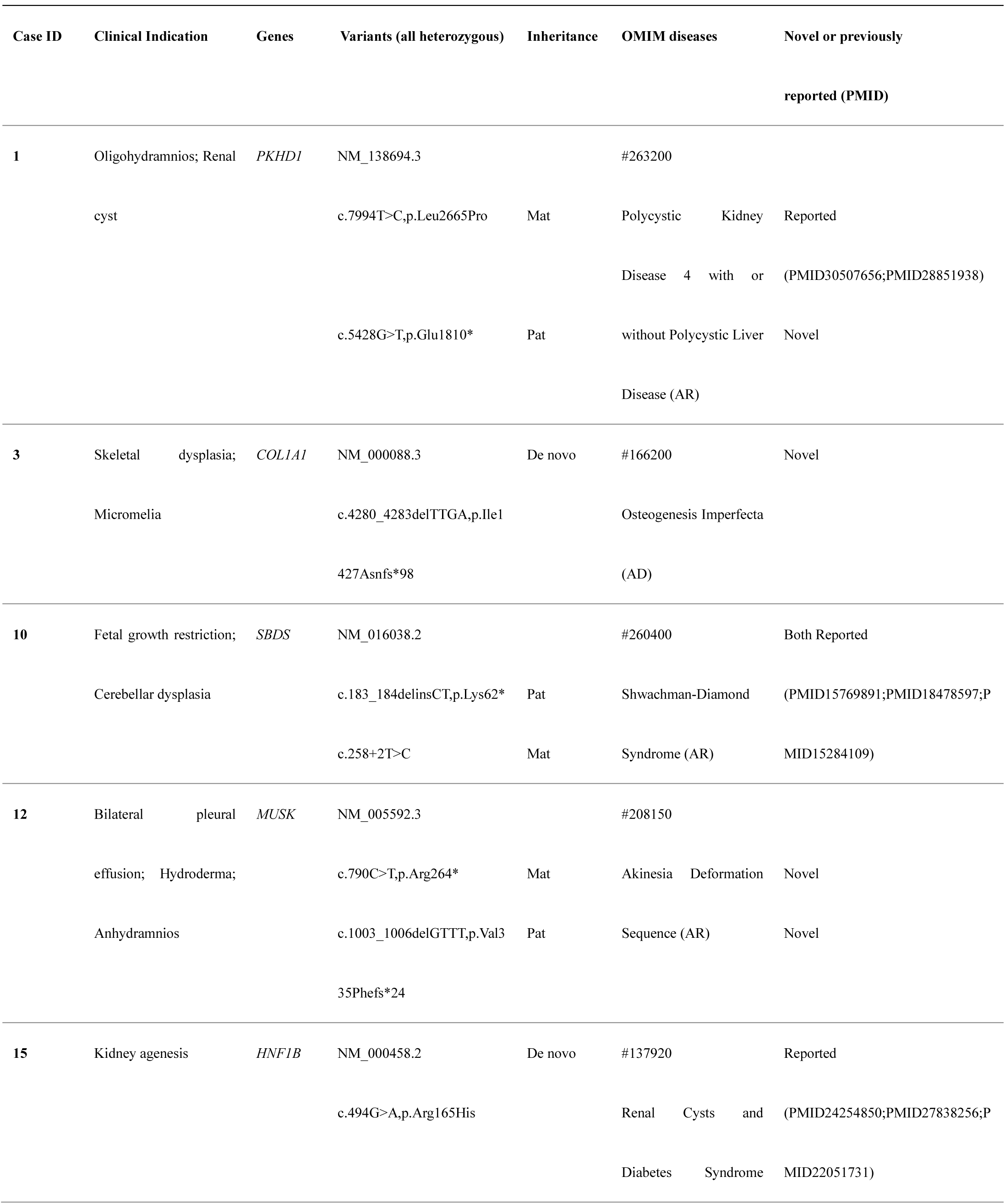
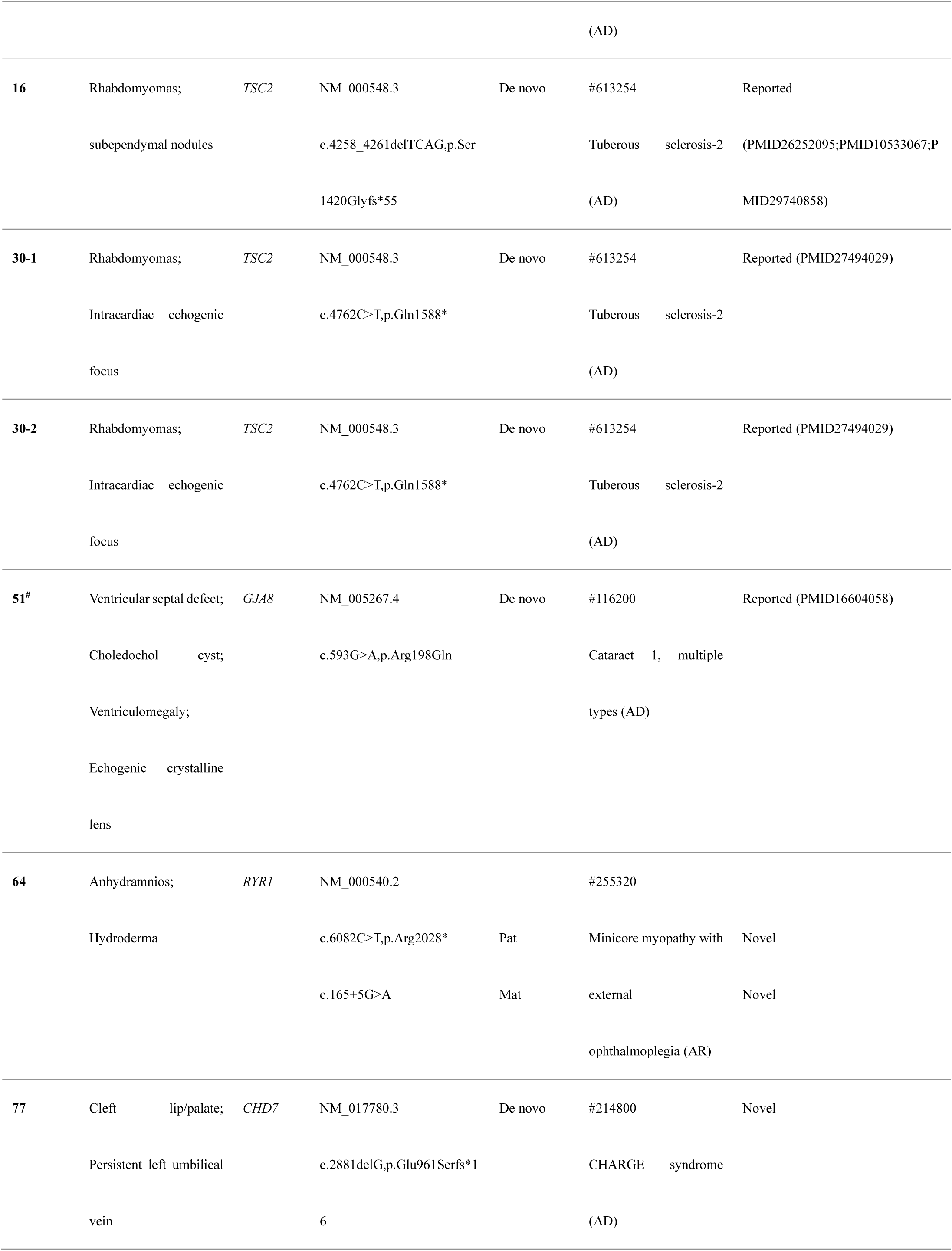
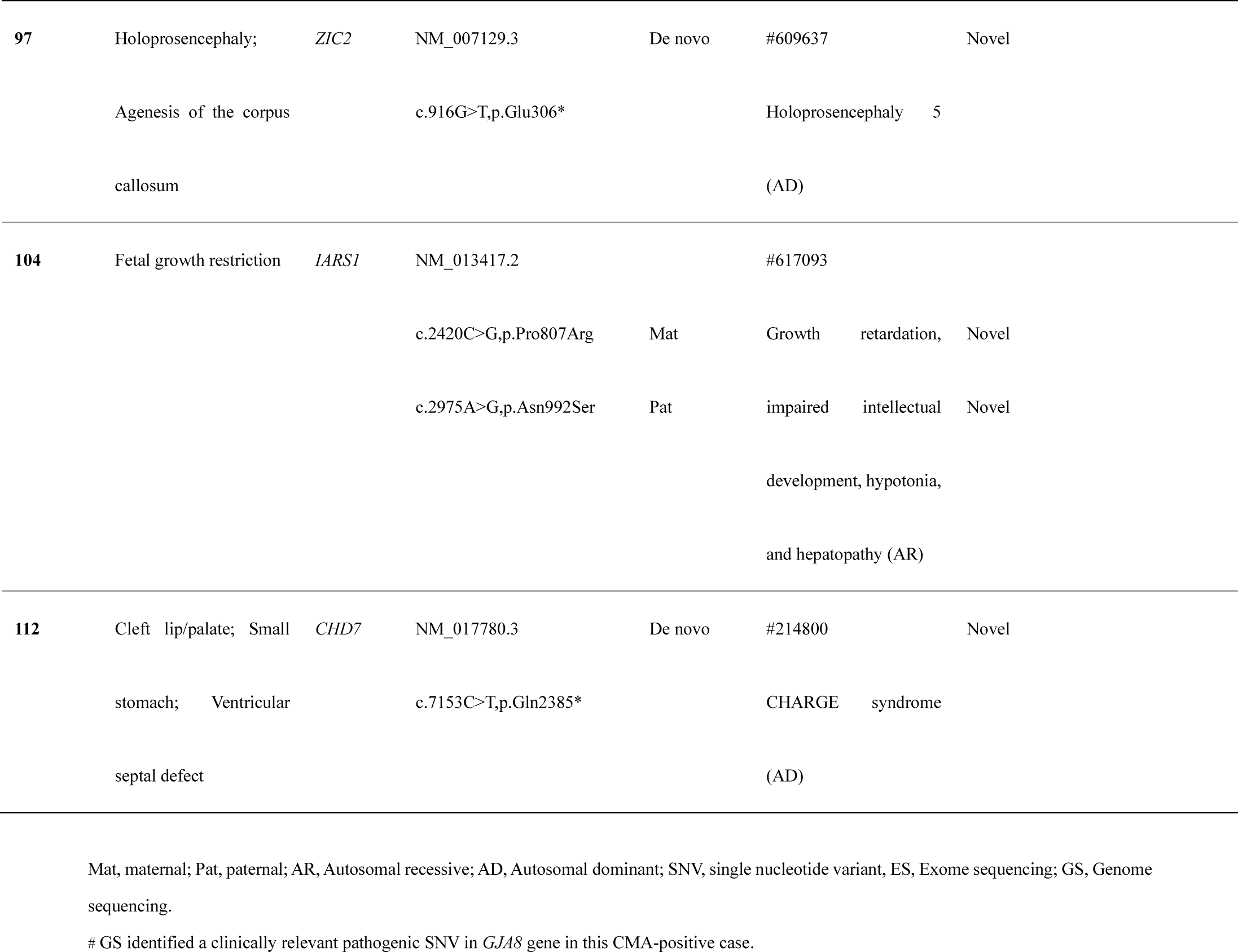
Summary of pathogenic or likely pathogenic single nucleotide variants (SNVs) and small insertions or deletions (INDELs) detected by ES and GS.

**Figure 2.**
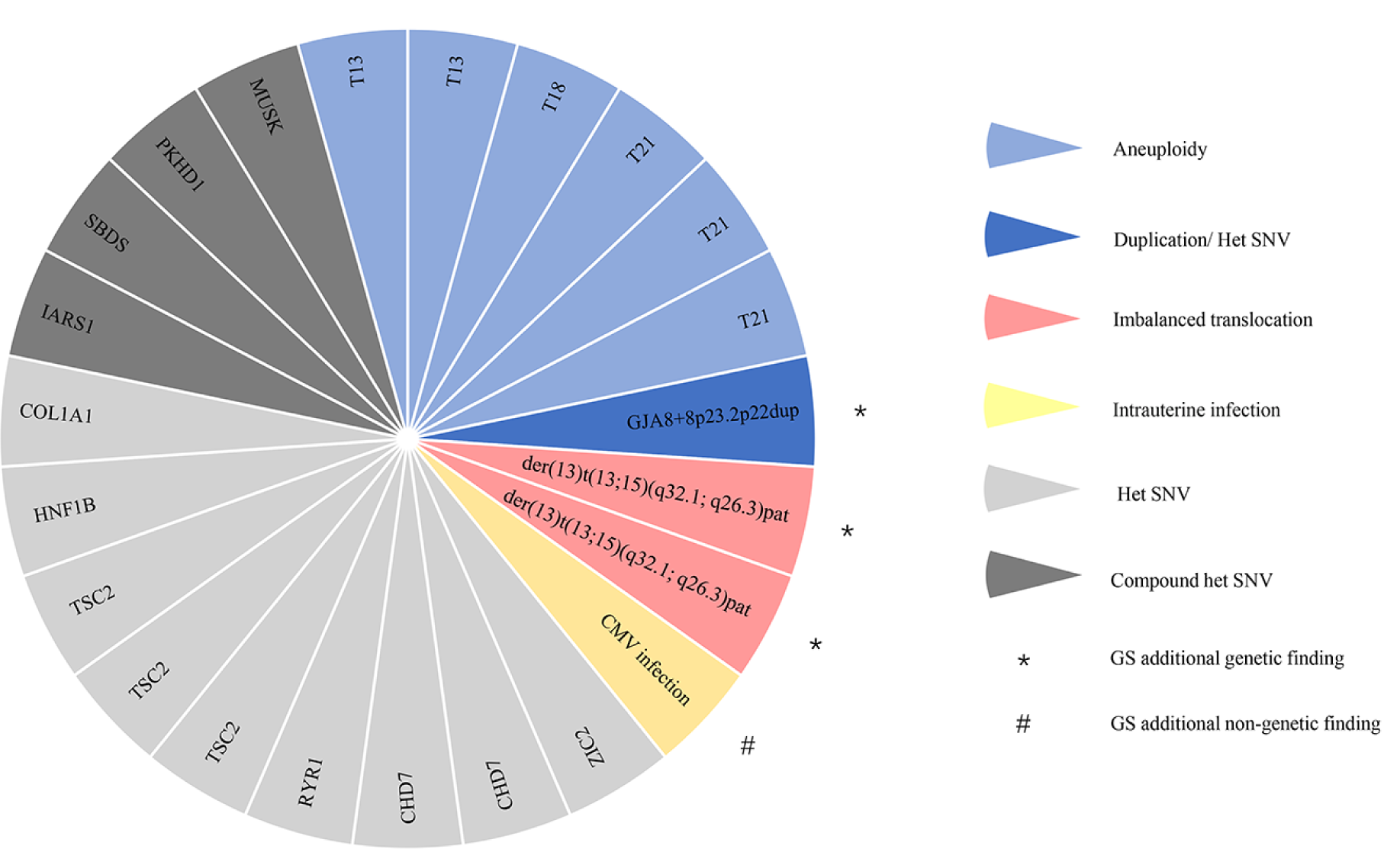
Architecture of a mixed cohort referred for prenatal diagnosis. Each slice of the pie chart represents one individual in the prospective cases analyzed by genome sequencing (GS) and chromosomal microarray (CMA) plus exome sequencing (ES) where clinically relevant findings were identified. Types of variant are indicated by colors (aneuploidy, light blue; duplication and heterozygous (het) single nucleotide variant (SNV), blue; imbalanced translocation, red; intrauterine infection, yellow; compound heterozygous SNV, light gray; heterozygous SNV, dark gray). Additional genetic findings by GS are indicated by * and additional nongenetic findings by GS are indicated by #.

**Figure 3.**
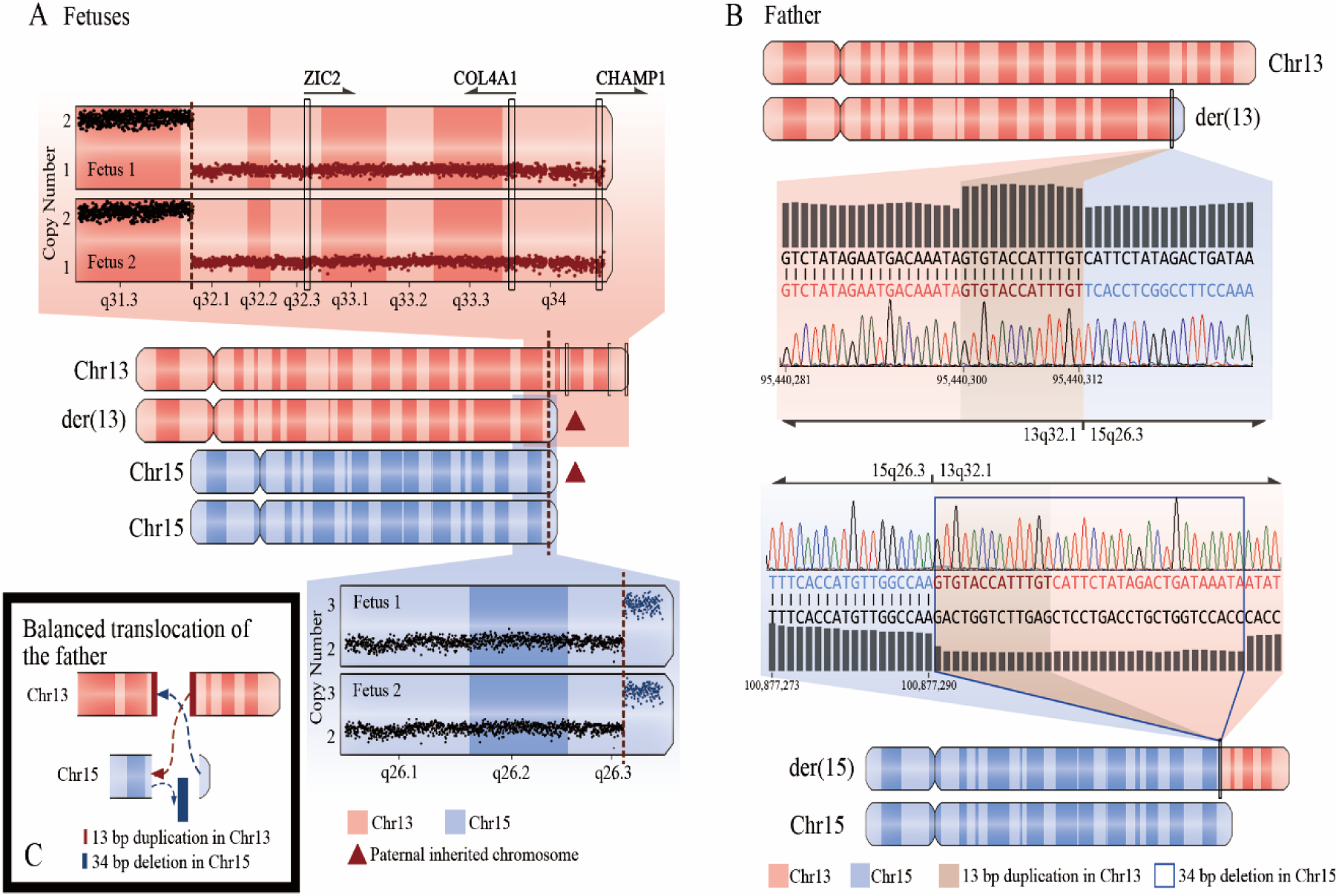
Genome sequencing (GS) identified pathogenic copy number variants (CNVs) in twin fetuses arising from their father’s balanced translocation. (**A**) Distributions of copy number across chromosome 13 and chromosome 15 are shown at the top and bottom, respectively. Dots in red indicate a heterozygous deletion of 19.7 Mb on chromosome 13, and dots in blue indicate a duplication of 1.6 Mb on chromosome 15 in both case 102-1 and case 102-2. The deletion affected 3 genes that are associated with autosomal dominant developmental disorder. (**B**) The father’s karyotype of chromosome 13 and chromosome 15 are shown at the top and bottom. The distribution of base pair sequencing depth and Sanger sequencing results across the breakpoints on chromosome 13 and chromosome 15 are displayed, respectively, in the middle. There is a templated 13-bp duplication and a 34-bp deletion in the breakpoint junction on chromosomes 13 and 15. (**C**) A schematic drawing of the formation of balanced translocations between chromosomes 13 and 15 in the father.

In addition, the potential for GS to provide a comprehensive picture of the molecular landscape for genetic interpretation was also reflected in the concurrent detection of CNVs and SNVs. For example, in case 51 with ventricular septal defect, choledochal cyst, ventriculomegaly and echogenic crystalline lens (suspected congenital cataract), 3 *de novo* pathogenic CNVs were detected by both CMA and GS. One of these CNVs, an ∼11.7-Mb duplication on chromosome 8p22-8p23.2 associated with 8p23.1 duplication syndrome[30], whose common features in the reported prenatal cases included congenital heart disease and whose novel features in the reported postnatal cases included ocular anomalies, was considered to explain the fetal phenotype; thus, ES was not carried out in this case due to the positive CMA finding. However, a clinically relevant *de novo* pathogenic SNV NM_005267.4, c.593G>A (p. Arg198Gln) in the *GJA8* gene, which was associated with Cataract 1, multiple types (OMIM: #116200), were identified by GS (Figure 2). This SNV has been reported in three children and their affected mother. All of them were diagnosed with bilateral cataracts soon after birth, and the variant detected in their mother via segregation analysis was *de novo*[31]. The SNV has also been reported in three affected members of an Indian family with a cataract history and was absent in 400 normal controls[32]. After comprehensive evaluation by clinicians and geneticists, both the SNV and the pathogenic CNV were considered to contribute to the fetus’s phenotype. However, the SNV was missed by the current CMA plus ES strategy.

To further demonstrate the capacity of GS to detect CNVs and SNVs/INDELs, we first compared all of the CNVs with sizes ≥100 kb detected between CMA and GS since the reporting threshold of CytoScan 750K we adopted for CMA was 100 kb. GS not only detected all of the CNVs with sizes ≥100 kb, which were detected by CMA, but also detected additional CNVs that CMA did not detect (Table S2). In addition, CNVs with sizes smaller than 100 kb were also detected by GS (Table S3), but none of these small CNVs were determined to be pathogenic. Our data indicated that GS was at least as powerful as CMA in the detection of CNVs with sizes ≥100 kb and had more potential to detect smaller CNVs. Second, we compared ES and GS in terms of their abilities to detect SNVs/INDELs in the coding region. Although the mean sequencing depth was higher in ES (147.1-fold [SD 24.0] vs. 57.6-fold [SD 4.2]), the coverage at 20-fold of GS was higher (97.8% [SD 1.3] vs. 96.7% [SD 1.3]), and GS detected more candidate variants in the coding region than ES (95% CI, p < 0.0001, paired t-test, Table S4), which indicated that GS is more powerful than ES for detecting exome variants.

### Additional findings of GS

With clinical GS, there is potential for recognition and reporting of incidental findings unrelated to the indication for ordering sequencing and even of the genetic information of associated microorganisms due to the noncapture method of GS. In case 5, a fetus with fetal growth restriction was incidentally found by GS to be infected with cytomegalovirus (CMV) (Figure 2), while the results of CMA and ES were both negative for CMV. This case was further validated by amplification of the viral DNA in the amniotic fluid through RT-PCR (Figure S2). CMV is the leading infectious cause of newborn malformation[33]. Inspired by this case, we developed a bioinformatics pipeline for the detection of pathogens associated with adverse perinatal outcomes, such as CMV, Toxoplasma (TOX), and Herpes simplex virus 1/2 (HSV 1/2), and analyzed all of the fetal samples; however, no additional positive cases were identified. Even so, GS showed its potential to provide information on nongenetic factors that neither CMA nor ES could provide in this case.

### Subgroup analysis

The 111 fetuses that were eligible for GS were categorized into 10 phenotypic groups based on the types of anomalies detected by ultrasound. The molecular diagnostic rate among 10 different phenotypic groups ranged from 0%-39.1% (9/23), as shown in Table S5. The highest diagnostic rate of 39.1% (9/23) was achieved for fetuses with multisystem anomalies, followed by 30.8% (4/13) for fetuses with cardiac anomalies. We did not find any diagnostic genetic variants in 5 fetuses with abdominal anomalies or in 7 fetuses with chest anomalies. Among 9 cases diagnosed with multisystem anomalies, 8 (88.9%) cases had chromosomal disorders, and 1 (11.1%) case had a monogenic disorder.

### Impact on pregnancy outcome

The effect of genetic diagnosis on pregnancy outcome is shown in Table 3. Follow-up results were available for all 111 fetuses. Among those of 22 diagnosed fetuses, the parents of 21 (95.5%, 21/22) fetuses opted for termination, and the parents of the remaining fetus (4.5%, 1/22) with clubhands, facial abnormality, omphalocele and umbilical cord cyst opted to continue the pregnancy. Among those of 89 undiagnosed fetuses, the parents of 34 (38.2%, 34/89) fetuses opted for termination, and the parents of 55 (61.8%, 55/89) fetuses opted to continue the pregnancies (Table S1). We noticed that there were no significant differences in the severities of abnormal fetal phenotypes between the diagnosed group and the undiagnosed group. Thus, in the context of an abnormal fetal phenotype, parents opted for termination of pregnancy significantly more often when they received a positive genetic diagnosis (p=0.00000053, Fisher’s exact test), which indicated that the specific genetic diagnosis influenced parental decision-making.

**Table 3.** Impact of genetic diagnoses on pregnancy outcome.

| Groups | Number of cases | Termination of pregnancy |  | Continuing pregnancy |  | P value<sup>^</sup> |
| --- | --- | --- | --- | --- | --- | --- |
|  |  | Number of cases | 95% CI<sup>#</sup> | Number of cases | 95% CI<sup>#</sup> |  |
| Diagnosed | 22 (19.8%) | 21 (95.5%) | 77.2%-99.9% | 1 (4.5%) | 0.1%-22.8% | 0.00000053 |
| Undiagnosed | 89 (80.2%) | 34 (38.2%) | 28.1%-49.1% | 55 (61.8%) | 50.9%-71.9% |  |
| Overall | 111 | 55 (49.6%) | 39.9%-59.2% | 56 (50.4%) | 40.8%-60.1% | - |

<sup>#</sup>95% confidence interval was calculated by binomial exact calculation.
<sup>^</sup>Fisher's exact test.

## DISCUSSION

In this prospective study, we applied GS in parallel with CMA plus ES to 111 fetuses with a broad range of structural or growth anomalies. To our knowledge, this is the largest prospective study on the use of high-coverage trio GS for the prenatal diagnosis of fetal structural abnormalities. GS, as a single test, revealed positive genetic findings in 22 fetuses, corresponding to a diagnostic rate of 19.8% (22 in 111). The same disease-causing variants were detected by GS and CMA plus ES in 19 fetuses of these fetuses, including 6 fetuses with chromosomal aneuploidies and 13 fetuses with P/LP SNVs or INDELs. For the remaining 3 fetuses, GS provided more comprehensive and precise genetic information than CMA plus ES, revealing twin fetuses with an imbalanced translocation and one fetus with an additional pathogenic variant in the *GJA8* gene. Furthermore, GS also incidentally identified intrauterine CMV infection in a growth-restricted fetus. Taken together, these findings show that GS can detect more types of variants and even provide clinically relevant nongenetic information that might not be detectable by CMA or ES, which indicates that GS has the potential to serve as a promising first-tier approach in prenatal diagnosis.

Previous studies[4, 6, 8, 11] have demonstrated the clinical utility of CMA and ES in prenatal diagnosis, and a combination of these two approaches for each case has been warranted. In our study, the capacity of GS for the prenatal diagnosis of fetal structural anomalies was more powerful than that of CMA plus ES in the following ways. First, precise breakpoint identification and analysis through GS data could detect more types of variants. In cases 102-1 and 102-2, twins with severe hydroderma and pleural effusion, an imbalanced translocation between chromosome 13 and chromosome 15 arising from a balanced paternal translocation was identified by GS, while CMA identified only one duplication and one deletion on these two chromosomes. The precise genetic diagnosis via GS allowed an accurate assessment of the recurrence risk in future pregnancies for this family. Second, GS can avoid the omission of the disease-causing variants by the clinical CMA plus ES sequential test. For example, in case 51, a *de novo* pathogenic SNV in the *GJA8* gene was identified by GS, while ES was not carried out due to the positive CMA results. GS provided more comprehensive genetic information in this case. Finally, GS was able to provide clues regarding the presence of nongenetic factors by incidentally identifying intrauterine CMV infection in a growth-restricted fetus. The pregnant woman was determined to be CMV-IgG positive and CMV-IgM negative before pregnancy, which indicated that her preconception immunity against CMV had been established. Given that the rate of fetal transmission in nonprimary infection is low, at 0.15-2%[34, 35], the intrauterine infection with CMV was not suspected at first; however, GS was able to provide the nongenetic information on an infection that had been overlooked in this case.

In addition, GS also demonstrated its advantages of a more rapid turnaround time (TAT) and lower input-DNA requirement in this study. The diagnostic process of CMA plus ES is time-consuming, with a median TAT of 10 (SD 2) days for CMA and 21 (SD 6) days for ES, and a total amount of ∼400 ng DNA would be required from limited prenatal samples. However, the GS approach requires a lower amount of fetal DNA (100 ng) and provides a more rapid median TAT of 18 (SD 6) days (Table S6), which is meaningful for such a time-sensitive analysis[36] since it reduces anxiety and stress for families and allows couples to have adequate time for genetic counseling and decision-making.

Of note, the prenatal detection rate for P/LP CNVs (2.7%) in our study was significantly lower than those in previous large-scale studies (4.3%-8.2%)[2, 11, 37]. This may be because the majority of fetuses (80.2%, 89 of 111 fetuses) in our study received serum screening or noninvasive prenatal screening (NIPS), and common P/LP CNVs were excluded before referral to our fetal medicine unit and prenatal diagnosis center. It is important to note that 6 fetuses with chromosomal aneuploidy underwent GS in our study. We speculated that there are two reasons for this. 1) The sonographic examination determined fetal abnormalities at an earlier gestational age before the serum screening and NIPS test, and these fetuses met our inclusion criteria and were included in our study cohort. 2) In resource-limited settings or due to the personal preference of the pregnant woman, serum screening and NIPS were not offered, and their fetuses were determined to be abnormal afterward. These cases were also included. Indeed, in our cohorts, 22 fetuses referred for CMA did not undergo serum screening or NIPS, and this resulted in 6 (27.3%) fetuses with trisomy 21/18/13 still being identified during GS implementation. We also note that the prenatal detection rate for P/LP SNVs/INDELs in our study was lower than that in a previous study of 50 fetuses with increased nuchal translucency (11.7% vs 14%)[18], possibly due to our larger cohort and the broader range of fetal structural anomalies in our study. However, the incremental diagnostic yield of ES in CMA-negative cases was higher in this study (12.7% vs 8.5%-10%)[8, 9] than in previous prospective studies on the prenatal application of ES in fetuses with structural or growth anomalies who had normal CMA results. One possible reason is the smaller cohort of this study.

However, the clinical usefulness of our GS scheme may be further improved, and many considerations should be addressed before implementing GS in clinical practice. First, we noticed that 6 fetuses with chromosomal aneuploidy went through our GS process. Thus, a reliable, rapid and cost-effective method, such as quantitative fluorescence polymerase chain reaction (QF-PCR), was needed to preliminarily exclude common fetal chromosomal aneuploidies before GS in clinical application, which could optimize the cost-effectiveness of the analysis[38]. Second, a large number of VUS variants in the noncoding region and incidental findings are called by GS, and a major concern is whether they should be reported prenatally. This decision is a particularly challenging because of the uncertainty associated with these findings, which further increases the complexity of genetic counseling compared to ES and increases difficulty in parental decision-making. Additionally, the choice to report the finding should also be weighed against the risk of missing a potential molecular diagnosis if not reported. Finally, even though sequencing costs continue to decrease, the cost of GS is comparatively high, especially for trio sequencing. Studies evaluating the cost-effectiveness of GS are warranted before clinical implementation.

In conclusion, our data show that GS provides comprehensive detection of various genomic variants in fetuses with structural or growth anomalies. In lieu of two separate analyses, GS, as a single test with a rapid TAT, should be performed as it has an equivalent diagnostic rate to that of CMA plus ES and provides a more comprehensive picture of the molecular landscape for the explanation of the mechanism. Although prospective studies with larger cohorts are warranted, our study demonstrates the feasibility of replacing CMA plus ES with GS as a first-line test for prenatal diagnosis in the near future.

## ACKNOWLEDGMENTS

We wish to extend our deepest gratitude to the individuals who participated in this research. Authors from Shanghai First Maternity and Infant Hospital designed the study, enrolled patients and collected samples, determined the detailed fetal clinical phenotype, performed CMA detection and interpretation, provided genetic counseling, followed up patients and revised the manuscript. BGI Genomics codesigned the study, performed GS and ES detection and interpretation, analyzed and summarized data, drafted the initial manuscript, and revised the manuscript. The final diagnosis was determined by a multidisciplinary team of fetal medicine specialists, genetic counselors and geneticists from the above cooperation units. This work was supported by a grant from the National Key Research and Development Program of China (No.2018YFC1002900). This work was also supported by China National GeneBank (CNGB). The data that support the findings of this study have been deposited into CNGB Sequence Archive [39] of CNGBdb [40] with accession number CNP0001235.

