## Supplementary figures and images for "Comprehensive genome sequencing analysis as a promising option in the prenatal diagnosis of fetal structural anomalies: a prospective study"

### Supplemental Figure 1

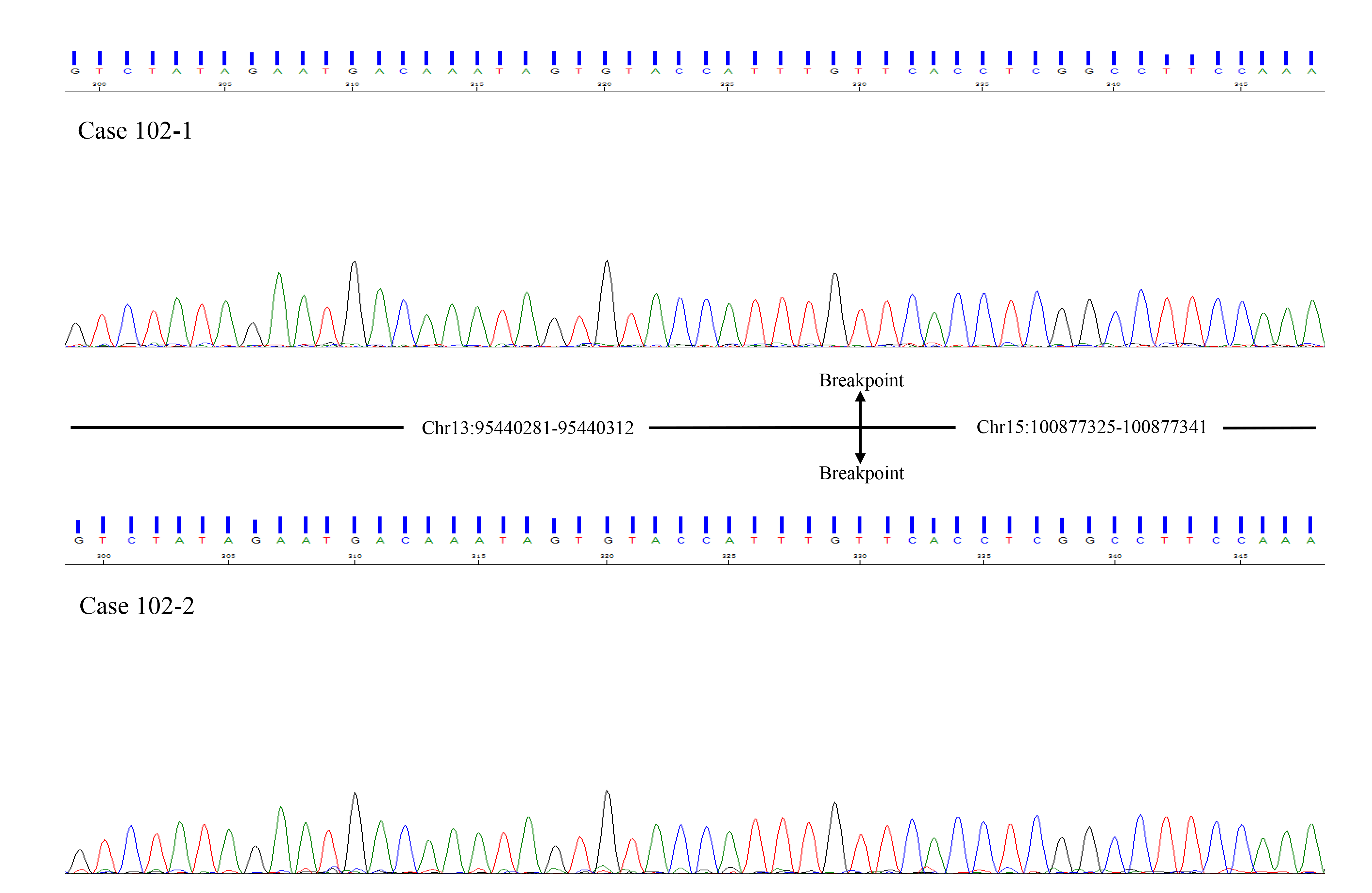

### Supplemental Figure 2

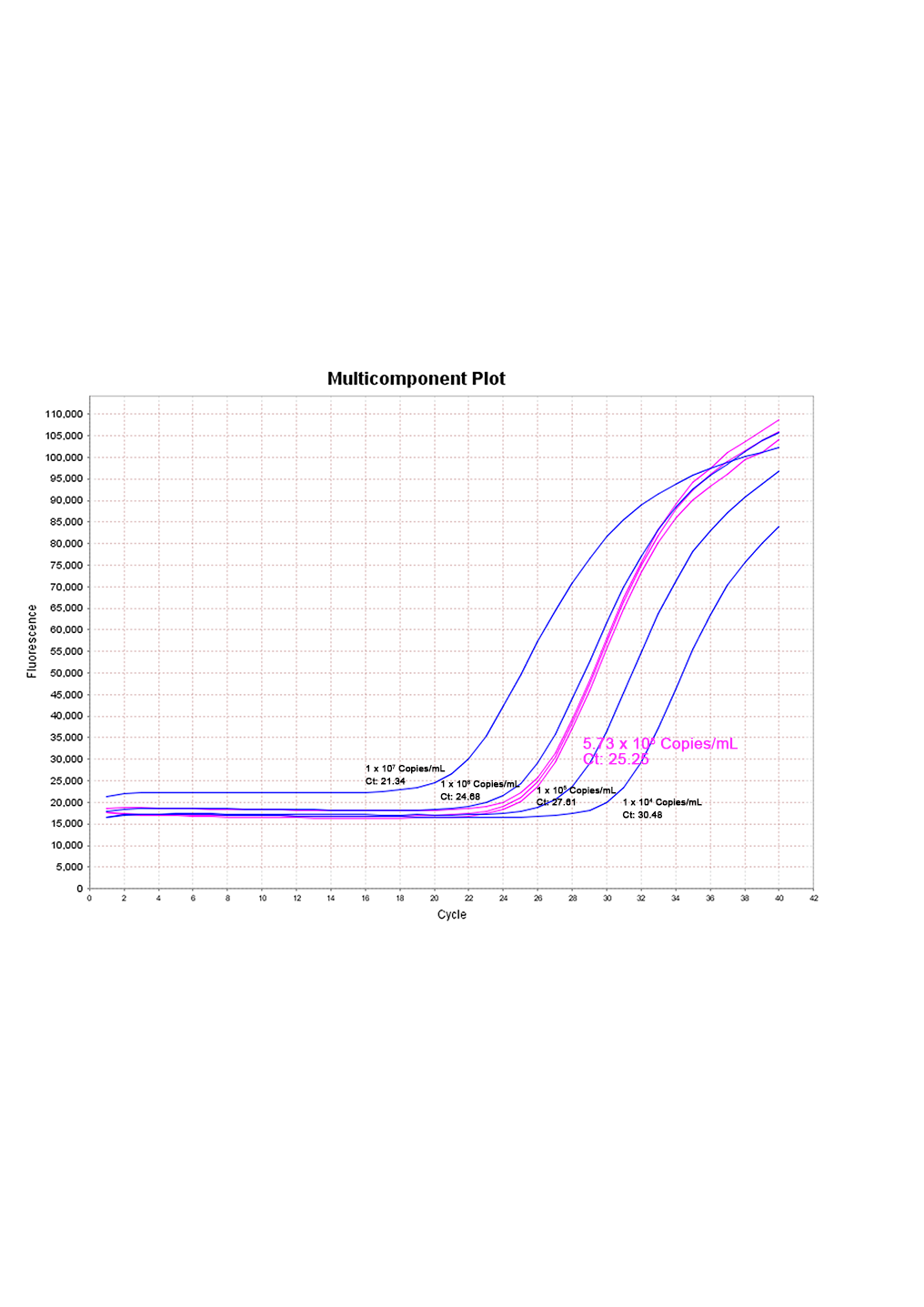
