## Supplementary materials and methods for "Comprehensive genome sequencing analysis as a promising option in the prenatal diagnosis of fetal structural anomalies: a prospective study"


#### Authors' full name and academic degrees:

Jia Zhou, MD[1], Ziying Yang, MS[2,3], Jun Sun, PHD[2,3], Lipei Liu, PHD[2,3], Xinyao Zhou, PHD[1], Fengxia Liu, MS[2,3], Ya Xing, MD[1], Shuge Cui, MS[2,3], Shiyi Xiong, MD, PHD[1], Xiaoyu Liu, MS[2,3], Yingjun Yang, MD[1], Xiuxiu Wei, MS[2,3], Gang Zou, MD[1], Zhonghua Wang, PHD[2,3], Xing Wei, MD[1], Yaoshen Wang, BS[2,3], Yun Zhang, MD[1], Saiying Yan, MS[2,3], Fengyu Wu, MD[1], Fanwei Zeng, MS[2,5], Tao Duan, MD[1], Jian Wang, PHD[4], Yaping Yang, PHD[6], Zhiyu Peng, PHD[2], Luming Sun, MD, PHD[1]

#### Authors' primary affiliations:

[1] Shanghai First Maternity and Infant Hospital, Tongji University School of Medicine, Shanghai, China;

[2] BGI Genomics, BGI-Shenzhen, Shenzhen, China;

[3] Tianjin Medical Laboratory, BGI-Tianjin, BGI-Shenzhen, Tianjin, China;

[4] Department of Medical Genetics and Molecular Diagnostic Laboratory, Shanghai Children's Medical Center, Shanghai Jiaotong University School of Medicine.

[5] Department of Biology, Faculty of Science, University of Copenhagen, Copenhagen, DK-2200, Denmark;

[6] AiLife Diagnostics, Pearland, TX 77584, USA.

These authors contributed equally: Jia Zhou, Ziying Yang and Jun Sun

1. Supplementary Methods

1-1. WGS single nucleotide polymorphisms (SNPs) / small insertions or deletions (INDELs) analysis

1-2. WGS copy number variants (CNVs) analysis

1-3. WGS structural variants (SVs) analysis

1-4. WGS runs of homozygosity (ROHs) analysis

1-5. WGS pathogens analysis

1-6. References for Supplementary Methods

2. Supplementary Figure Legends

### 1. Supplementary Methods

#### 1.1 WGS SNPs/INDELs analysis

In this part, all the preprocessing steps of data from raw reads to final bam are implicated, then do all kinds of analysis.

##### 1.1.1 Filter

Low quality reads deleting and removed for next analysis[1].

```
fastp --thread 8 –I [fastq file for the first of read pairs] –I [fastq file for the second of read pairs] –o [cleaned fastq file for the first of read pairs] –O [cleaned fastq file for the second of read pairs] -s 10 –j [statastics file in json format ] –h [statastics file in html format ]
```

##### 1.1.2 Bam

Aligning clean reads to human reference genome, sorting by coordinate, marking duplication reads for PCR duplication, fixing mate information, and detecting and correcting of systematic errors[2-5].

1) Align.

Alignment with clean reads

```
bwa mem -M -t 8 -Y –R "@RG\tID:sample id\tSM:sample id\tPL:platform" [GRCh37 reference] [cleaned fastq file for the first of read pairs]  [cleaned fastq file for the second of read pairs] | samtools view -S -b –o [output BAM file] -
```

2) Sort

```
sambamba sort --memory-limit 23G -l 1 -t 8 --tmpdir=[temporary output directory] –o [output aligned BAM] [input BAM]
```

3) Merge

```
samtools merge -R [chromosome tag] -c -p –f [output merged bam] [all bams of parts reads aligned ]
```

4) Dup

```
java -Xmx20G -XX:ParallelGCThreads=2 \
-Djava.io.tmpdir=[temporary directory] –jar MarkDuplicates.jar INPUT=[each chromosome bam] OUTPUT=[output bam file] METRICS_FILE=[statistical file]
VALIDATION_STRINGENCY=SILENT
MAX_FILE_HANDLES_FOR_READ_ENDS_MAP=8000
```

5) Fix

```
java -Xmx20G    -Djava.io.tmpdir=[temporary directory]    –jar gatk.jar FixMateInformation
--VALIDATION_STRINGENCY SILENT –I [input bam] -O [output bam]
```

6) Bqsr

```
java -jar gatk.jar BaseRecalibrator –R [GRCh37 reference] –I [input bam] --tmp-dir [temporary directory] --known-sites [1000G vcf file] --known-sites [1000genome snp vcf file] --known-sites [dbSNP vcf file] --known-sites [1000genome snp vcf file] –O [1000genome snp vcf file]

java -Djava.io.tmpdir=[tmp directory] –jar gatk.jar ApplyBQSR –I [input bam] –bqsr [bqsr mediate file] –O [output bam file]
```

### 1.1.3 QC

Statistics about the depth, coverage, Q20, Q30, GC content and so on

1) Bam split

```
Sambamba view -f bam -h -o [output bam] -L [bed] [input bam]
```

2) Partial bam qc

```
bamdst -p [bed] -o [output directory] [input bam]
```

3) QC collection

```
Python wgs.qc_collect.py –filter [filter directory] –bamqc [ partial bam qc ] -o [output directory] --sex [sex tag] > [output file]
```

1.1.4 SNPs/INDELs analysis

Using GATK HaplotypeCaller model, generating SNPs and INDELs saved as VCF file, and also the gvcf file, then going on VQSR step reducing the corrected quality of the variation.

1) Variant calling

```
java -Xmx4G -XX:ParallelGCThreads=4 –jar gatk.jar HaplotypeCaller --tmp-dir tmp -ERC GVCF --correct-overlapping-quality true -A BaseQuality -A MappingQuality -A QualByDepth -A MappingQualityRankSumTest -A ReadPosRankSumTest -A FisherStrand -A StrandOddsRatio -A InbreedingCoeff –R [reference file] –L [bed] –I [input bam] –O [output gvcf file]

java -Xmx4G -XX:ParallelGCThreads=4 –jar gatk.jar GenotypeGVCFs --tmp-dir tmp –R [reference file] –V [gvcf file] –L [bed] –O [vcf file]
```

2) Variant collection

```
bcftools concat -a -D -q 30 -O z –o [output vcf file] –f [vcf list file]

tabix -p vcf [input vcf]
```

3) SNPs vqsr

```
java –jar gatk.jar SelectVariants --tmp-dir=snp_javatmp –R [reference file] –variant [vcf file] –O [snp vcf file] -select-type SNP

java –jar gatk.jar VariantRecalibrator --tmp-dir=snp_javatmp –R [reference file] –V [snp vcf file] --resource hapmap,known=false,training=true,truth=true,prior=15.0:hapmap_3.3.hg19.sites.vcf.gz --resource omni,known=false,training=true,truth=false,prior=12.0: 1000G_omni2.5.hg19.sites.vcf.gz --resource 1000G,known=false,training=true,truth=false,prior=10.0: 1000G_phase1.snps.high_confidence.hg19.sites.vcf.gz --resource dbsnp,known=true,training=false,truth=false,prior=2.0:dbsnp_138.hg19.vcf.gz -an DP -an QD -an MQ -an MQRankSum -an ReadPosRankSum -an FS -an SOR -mode SNP –O [temporary vqsr file of snp vcf ] --tranches-file [snp tranches vcf file]

java –jar gatk-package-4.0.11.0-local.jar ApplyVQSR --tmp-dir=[tmp directory] –R [reference file] –V [snp vcf file] –O [output snp vcf file] --truth-sensitivity-filter-level 99.0 --tranches-file [snp tranches vcf file] --recal-file [temporary vqsr file of snp vcf ] -mode SNP

java –jar gatk.jar SortVcf –I [filtered snp vcf file] –O [output vcf file]
```

4) INDELs vqsr

```
java –jar gatk.jar SelectVariants --tmp-dir=[temporary directory] –R [reference file] –[input vcf file] –O [output snp vcf file] -select-type INDEL

java –jar gatk.jar VariantRecalibrator --tmp-dir= indel_javatmp –R [reference file] –V [indel vcf file]-resource mills,known=true,training=true,truth=true,prior=12.0: Mills_and_1000G_gold_standard.indels.hg19.sites.vcf -an DP -an QD -an MQ -an MQRankSum -an ReadPosRankSum -an FS -an SOR -mode INDEL –O [temporary vqsr file of indel vcf] --tranches-file [indel tranches vcf file]

java –jar gatk.jar ApplyVQSR --tmp-dir= [temporary directory] –R [reference file] –V [indel vcf file] –O [output indel vcf file] --truth-sensitivity-filter-level 99.0 --tranches-file [indel tranches vcf file] --recal-file [temporary vqsr file of indel vcf ] -mode INDEL

java –jar gatk.jar SortVcf –I [filtered indel vcf file] –O [output vcf file]
```

5) SNPs/INDELs concat

```
java –jar gatk.jar MergeVcfs –I [input snp vcf file] –I [input indel vcf file] –O [output vcf file]

tabix -p vcf –f [input vcf file ]
```

#### 1.2 WGS CNVs analysis

CNVnator[6] used for the whole genome large CNV detection, and ExomeDepth[7] for small CNV detection.

1) CNVnator

```
chr=$1

root=$2

bam=$3

ref_dir=$4

out=$5

bin=$6

cnvnator=$7

$cnvnator -root $root -chrom $chr -tree $bam -unique

$cnvnator -root $root -his $bin -d $ref_dir

$cnvnator -root $root -stat $bin

$cnvnator -root $root -partition $bin

$cnvnator -root $root -call $bin >$out
```

2) ExomeDepth

```
#!/bin/bash

sampleID=$1

inbam=$2

gender=$3

outdir=$4



bin_dir=$(dirname $0)

CNV_anno=$bin_dir/CNV_anno
```

```
sourceDir=$bin_dir

bed=$sourceDir/hg19.chr.bed

Rscript=$bin_dir/../../tools/Rscript-exondepth

samtools=$bin_dir/../../tools/samtools

control1=$sourceDir/control/wgs_control/lfx/all.A.my.count.rds

control2=$sourceDir/control/wgs_control/lfx/all.$gender.my.count.rds

prefix=$(basename $inbam)

bam=$outdir/$prefix.hg19.chr.bed.bam

$samtools view -b -L $bed -o $bam $inbam

$samtools index $bam

script=$sourceDir/run.getBamCount.R

$Rscript $script $sampleID $bam A $outdir $sourceDir

echo `date` $Rscript $script $sampleID $bam $gender $outdir $sourceDir

$Rscript $script $sampleID $bam $gender $outdir $sourceDir

script=$sourceDir/run.getCNVsFromControl.R

echo `date` $Rscript $script $sampleID A $outdir $control1

$Rscript $script $sampleID A $outdir $control1

$Rscript $script $sampleID $gender $outdir $control2

echo $sampleID.A.CNV.calls.tsv > $outdir/all.CNV.calls.list

echo $sampleID.$gender.CNV.calls.tsv >> $outdir/all.CNV.calls.list

echo -e "$sampleID\t$gender" > $outdir/sample.list.checked
```

```
perl \
$CNV_anno/script/add_cn_split_gene.batch.pl \
$outdir/all.CNV.calls.list \
$outdir/sample.list.checked \
$CNV_anno/database/database.gene.list.NM \
$CNV_anno/database/gene_exon.bed \
$CNV_anno/database/OMIM/OMIM.xls \
$outdir/all.CNV.calls.anno.withoutHGMD


perl \
$CNV_anno/script/add_HGMD_gross.pl \
$outdir/all.CNV.calls.anno.withoutHGMD \
$CNV_anno/database/hgmd-gross_all-ex1-20190426.tsv \
$outdir/all.CNV.calls.anno
```

##### 1.3 WGS SVs analysis

SVs detection by software LUMPY[8].

1) Extract the discordant paired-end alignments

```
samtools view -b -F 1294 [input bam] |>[output bam]
```

2) Extract the split-read alignments

```
samtools view -b -F 1294 [input bam] extractSplitReads_BwaMem -i stdin|samtools view -
Sb - >[output bam]
```

3) SVs detection

```
lumpyexpress -B [input bam] -S [split bam] -D [disco bam] -T [temporary directory] -x [bed] -o [output vcf file]
```

##### 1.4 WGS ROH analysis

Runs of homozygosity detection by ROH software[9]

```
python -m  loh –vcf [filter vcf file] --wgs --out  [output directory] --id [sample ID]
```

##### 1.5 WGS pathogens analysis

Intrauterine infection statistics step by step.

```
mkdir -p fq
samtools fastq -@ 5 -f 4 [input bam] >[unmap fq]
mkdir -p bwa
bwa mem -R "@RG\tID:[sample ID]\tSM: [sample ID]\tLB:[library]\tPL:BGISEQ"  -t 5 -a -M [infection reference ] [unmap fq] | samtools view -S -b -o [raw bam]
samtools sort -o [sorted bam] [raw bam]
samtools index [sorted bam]
java -Djava.io.tmpdir=[temporary directory] -jar picard.jar MarkDuplicates MAX_FILE_HANDLES_FOR_READ_ENDS_MAP=8000 INPUT=[sorted bam] OUTPUT=[mark dup bam] METRICS_FILE=[dup.metrics ] VALIDATION_STRINGENCY=SILENT
samtools index [mark dup bam]
samtools flagstat  [mark dup bam]  >[stat file]
mkdir -p coverage
```

```
samtools view -h -F 4 -b [mark dup bam] -o [map bam]

samtools depth    [map bam] >[map bam depth]

perl depth2coverage.pl [map bam depth] [infection reference fai] [coverage]

samtools view -h -F 1028 -b [mark dup bam] -o [map and deldup bam]

samtools depth [map and deldup bam] >[map and deldup depth]

perl depth2coverage.pl [map and deldup depth] [map and deldup coverage]

perl cal_abu_by_reads_v1.pl [stat file] [database data] [coverage] [map bam]

[markdup.species_abundance file]

perl cal_abu_by_reads_v1.pl [stat file] [database data]    [markdup.species_abundance file]

[map and deldup bam] [markdup.species_abundance file 2]
```

#### 2 Supplementary Figure Legends

Figure S1. Sanger validation of the imbalanced translocation detected in twin fetuses. The derived chromosome 13 consists of chromosome 13 and chromosome 15 with the breakpoint located on chr13: 95440312.

Figure S2. RT-PCR validation of the viral DNA in the amniotic fluid. The load of cytomegalovirus (CMV) was 5.73 x 105 copies/mL, which indicated intrauterine CMV infection in case 5.
